# A Guide to Analysis and Reconstruction of Serial Block Face Scanning Electron Microscopy Data

**DOI:** 10.1101/133231

**Authors:** E. Cocks, M. Taggart, F.C. Rind, K. White

**Affiliations:** Newcastle University, Institute of Genetic Medicine, Central Parkway, Newcastle upon Tyne, NE1 3BZ

**Keywords:** Serial Block Face Scanning Electron Microscopy, Image Analysis, Skeletal Muscle, Fiji, Blender, Amira, Microscopy Image Browser

## Abstract

Serial block face scanning electron microscopy (SBF-SEM) is a relatively new technique that allows the acquisition of serially sectioned, imaged and digitally aligned ultrastructural data. There is a wealth of information that can be obtained from the resulting image stacks but this presents a new challenge for researchers - how to computationally analyse and make best use of the large data sets produced. One approach is to reconstruct structures and features of interest in 3D. However the software programs can appear overwhelming, time consuming and not intuitive for those new to image analysis. There are a limited number of published articles that provide sufficient detail on how to do this type of reconstruction. Therefore the aim of this paper is to provide a detailed step-by-step protocol, videos and explanation on the types of analysis and programs that can be used. To showcase the programs, skeletal muscle from fetal and adult guinea pigs were used. The tissue was processed using the heavy metal protocol developed specifically for SBFSEM. Trimmed resin blocks were placed into a Zeiss Sigma SEM incorporating the Gatan 3View and the resulting image stacks were analysed in 3 different programs, Fiji, Amira and MIB, using a range of tools available for segmentation. The results from the image analysis comparison show that the analysis tools are often more suited to a type of structure. For example larger structures, such as nuclei and cells, can be segmented using interpolation, which speeds up analysis; single contrast structures, such as the nucleolus, can be segmented using the contrast-based thresholding tools. Knowing the nature of the tissue and its specific structures (complexity, contrast, if there are distinct membranes, size) will help to determine the best method for reconstruction and thus maximising output from valuable tissue.

## Introduction

Electron Microscopy (EM) has evolved to incorporate different preparation techniques for the imaging of a wide variety of samples. Transmission and scanning EM (TEM and SEM respectively) are regularly used to analyse biological material in order to reveal structural information, which may relate to function. Yet, there are limitations. For example, SEM only images the surface topography of cells and tissues but not intracellular structures. On the other hand, TEM provides information on spatial arrangements within cells to within 1nm resolution. However, this is accomplished by examining a single ultrathin section, typically 50-100nm, from a much larger sample, which could range from ~10μm (single cells) to several mm (tissues). Therefore important information regarding spatial arrangements of structures of interest through the depth of a cell or tissue is difficult to obtain. Manual serial-sectioning can be combined with TEM to create 3D stacks and visualisations of data (Andersson-Cedergren, 1959; Harris and Stevens, 1988; Bock et al, 2011; Takemura, 2015 and Lee et al, 2016). However this is extremely time consuming, requires a high level of experience in microtomy and manual image alignment and, even then, often results in damaged or lost sections whose ‘missing’ information has to be interpolated.

The desire within the EM community to obtain data in three spatial dimensions (x-y-z) at the ultrastructural level led to the development of a rudimentary automated system for serial sectioning coupled with electron microscopy imaging by Kuzirian and Leighton in 1983. Although it was another two decades before the procedure known as Serial Block Face SEM (SBF-SEM) was described in published form by Denk and Horstmann (2004). With subsequent refinements of machinery and computational power, SBFSEM now fills many of the gaps in functionality of SEM and TEM. It consists of a mini-ultramicrotome with diamond knife inside an SEM chamber. The knife cuts ultrathin sections from a piece of tissue embedded in a resin block, an electron beam scans the block surface and a detector records the back-scattered electrons, producing a digital image. This process is repeated at an operator-specified depth to produce a digitised stack of aligned images. It is possible to obtain tens to hundreds of serial sections from resin blocks, and aligned images, in a few hours.

The stack of images obtained from biological samples allows researchers to follow cell-to-cell arrangements, or intracellular structures, in the z-axis in a number of ways. First, this can be achieved by simply scrolling through the images for qualitative assessment of the features of interest. Second, image analysis software can be used to create 3D reconstructions of the data. These can aid the qualitative assessment of the data, for example creating movies that show the reconstructions on rotating axis (Kasthuri et al, 2015). Third, such tools can also be used for detailed quantification of the biological data. The image analysis can be the most complex and time consuming portion of the whole process, especially for researchers new to electron microscopy and/or SBF-SEM.

In many ways the challenges of 3D EM have now shifted from how to capture the difficult-to-measure to what to do with all this data? At the outset of an experiment one needs to know how the resultant images are to be analysed. These considerations vary from simple to complex, depending upon the experimental question, the tissue or cell constituency, the resolutions of structures of interest and their contrast to neighbouring structures.

The most likely requirement is the creation of 3D reconstructions of the image stacks, whether for qualitative or quantitative assessment. There are a number of image data analysis packages that assist with creating reconstructions (for a recent review of these see Borrett and Hughes, 2016), each requiring segmentation as a first step. Segmentation is the process of annotating a specific structure on each image so as to follow it in each consecutive image in the z-axis. There are two main ways of accomplishing this. In one case, the object(s) is highlighted manually by adding a coloured layer(s) on top of the image. An alternative process involves assigning individual image pixels to only one object, this means if a pixel is reselected during another segmentation it will be reassigned.

The methods of segmentation can be further divided into manual, semi-automated and automated categories. Manual segmentation tools require the user to annotate the object, e.g. by colouring, over every slice. Semi-automated tools use a combination of user input and program predictions, to highlight a structure. An example of this is interpolation. The user manually annotates the structure every nth slice and the program will fill in the empty slices using the annotated image as a guide to predict the possible shape of the object. Another is the thresholding tool, which selects pixels based on the contrast limits set by the user. These limits allow for the selection of light or dark pixels depending on the appearance of the object. There is the option available in some programs for automated segmentation. Here, the program ‘learns’ object selection based on trial runs performed by the user on a sample dataset. These settings are then automatically applied to the full dataset to be analysed.

The complexity of SBF-SEM datasets, and scenario-specific analysis requirements, can make it difficult to know which image analysis tools and procedures to follow to make best interpretations of the data. Therefore, the purpose of this paper is to compare three popular image analysis programs, and the segmentations tools they offer, for the qualitative and quantitative analysis of SBF-SEM data. We provide detailed protocols for the data handling and analyses. In doing so, we aim to provide direction to researchers new to SBF-SEM by drawing attention to advantages and limitations of the software packages and tools. The material analysed by EM was guinea pig skeletal muscle, chosen because it exhibits regular, well-defined intracellular structures by EM that allowed us to test the effectiveness of each software tool. From the results of the analysis, we have developed a decision-making map that can be applied to the analysis of any structure from any sample type and will help researchers choose the workflow to apply to their SBF-SEM data. By following this workflow, the researcher will ensure that they are in the best position to address the aims of their study and maximise the output from this relatively new technique.

## Method

### Tissue preparation for SBF-SEM

The samples used here are skeletal tissue (psoas and soleus muscles) from Duncan Hartley guinea pigs terminated under licensed procedures according to the Animals Scientific Procedures Act 1986 (ASPA). Tissue was micro-dissected into 2% glutaraldehyde with 0.1M sodium cacodylate buffer and left for a minimum of 12hrs in the fixative. The samples then underwent a heavy metal staining protocol adapted from Deerinck (2010). The tissues were washed in 0.1M sodium cacodylate pH7.4 and then a solution of 3% potassium ferrocyanide with 2% aqueous osmium tetroxide in ddH2O is added. This is followed by filtered 10% thiocarbohydrazide, TCH and the secondary 2% osmium tetroxide. The samples are then placed in 1% uranyl acetate at 4oC overnight and then in lead aspartate solution, 0.12gms of lead nitrate in 20mls aspartic acid. Between each step the samples are washed in several changes of ddH2O. The samples are dehydrated with acetone, from 25% to 100% and then impregnated with increasing concentrations of Taab 812 hard resin in acetone with several changes of 100% resin. The samples are embedded into 100% resin and left to polymerise at 60°C for a minimum of 36 hours.

The resin blocks were trimmed, using a razor blade to form a trapezoid block face. Using a diamond knife, 1μm sections were taken and stained with toluidine blue and viewed under a light microscope. If tissue morphology and orientation was acceptable, several 70nm sections were taken, placed on a copper grid and viewed on a TEM. This is done to ensure that the tissue is adequately processed for viewing. The blocks were then further trimmed to approximately 0.75mm by 0.75mm and glued onto a pin. In order to reduce sample charging within the SEM, the block was painted in silver Dag and sputter-coated with a 10nm layer of gold.

### SBF-SEM settings

The specimens were placed into a Zeiss Sigma SEM (Zeiss, Cambridge, UK) incorporating the Gatan 3view (Gatan inc. Abingdon, UK) as the SBF-SEM system. For this particular project the following parameters were used. For each sample, the images were obtained at 4 or 5kv accelerating voltage, with an aperture of 30μm, in variable pressure ranging from 40 to 53Pa. The blocks were sectioned at a thickness of 70nm and the images recorded at a range of magnifications with a resolutions of 1024 by 1024 or 3000 by 3000 pixels with a 20μm/s dwell time.

Gatan Digital Micrograph was used to collect digitised images of each experimental run in a DM3 format. The data were then analysed using three different image analysis programs (Figure 1). These were Fiji (http://fiji.sc/), Amira (http://www.fei.com/software/amira-3d-for-life-sciences/) and Microscopy Image Browser, MIB, (http://mib.helsinki.fi/). Fiji, via the plugin TrakEM2 (Cardona et al., 2012), is primarily an operator-driven program that is freely-available. Amira and MIB (Belevich et al., 2016) have the capacity for using semiautomated tools: Amira requires a commercial license and MB is freely available. Amira enables visualisation of all analysed data, whereas FIJI and MIB are purely analytical programs and require a secondary program to visualise the segmentations. Blender (https://www.blender.org/) is a free graphics software which can be used to reconstruct the objects created in Fiji with the aid of Neuromoph Tools (Jorstad et al, 2014) (http://cvlab.epfl.ch/NeuroMorph) developed specifically to import the objects exported from Fiji and perform quantification analysis. Opening the segmented 3D objects via these tools also ensures that the dimensions are consistent with the parameters from the raw data. In MIB the model file can be exported in a variety of formats for different programs.

**Figure 1.**
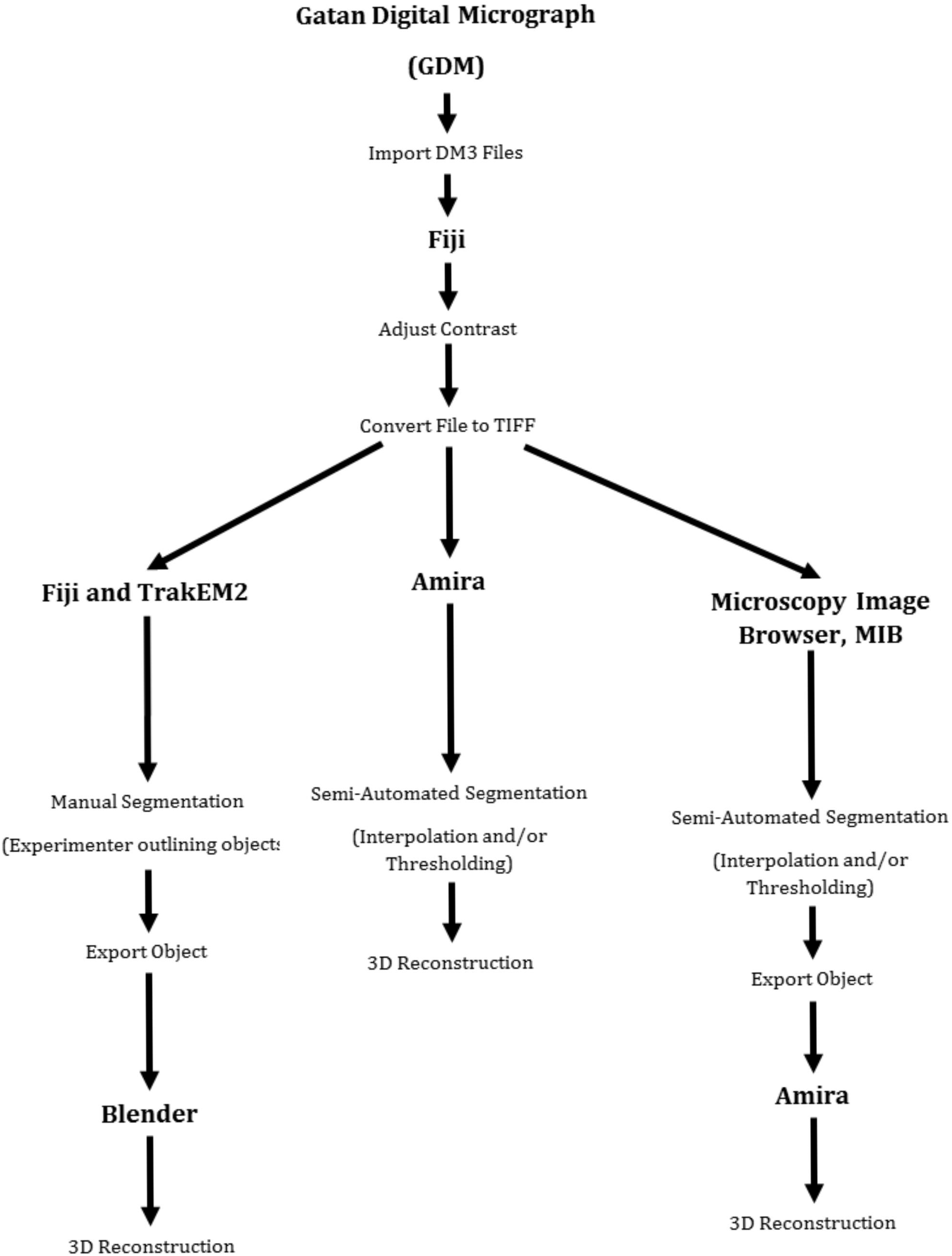
Flow Chart Showing the Steps for the Image Analysis Comparison. The first step was to adjust the contrast and convert the images from DM files formats to TIFF. The stack of TIFFs were then analysed in each of the 3 programs, the examples of the segmentations used in each are also shown. The final step in the process was to reconstruct the segmentations into a 3D model.

### Protocols for image file handling and analyses

The protocols below were applied to the examination of a skeletal muscle SBF-SEM dataset and the steps in each process illustrated in accompanying video file tutorials (Supporting files S1, S2, S3, S4 and S5). The dataset used in the videos can be accessed via the EMPIAR website, http://www.ebi.ac.uk/pdbe/emdb/empiar/, (Accession code: EMPIAR-10092).The protocols specific time points are given at the end of important steps, to aid in following and understanding how the programs work.

#### Converting DM3 to TIFF (Video Fiji Image Processing)

The raw data is saved as a DM3 file format from Gatan Digital Micrograph, GDM, which cannot be opened in all imaging analysis programs. So the first step was to convert the images into a TIFF format, during this process contrast can be lost so the contrast is normalised to ensure this does not happen during conversion.

1. Image sequence is imported by selecting ‘File’ > ‘Import’ > ‘Image Sequence’. *(see from time 00:06 in supplementary video file S1)*
2. The first image, or entire folder of pertinent images, is selected. A new window appears and the number of images to be imported is checked. Ensure that nothing else is ticked.
3. The image stack will then open.
4. The contrast is normalised by choosing ‘Process’ > ‘Enhance Contrast’. *(time 00:50, file S1)*
5. Saturated pixels are changed to 1% and ‘Normalise’ and ‘Process All’ are ticked.
6. Contrast can be further enhanced by going to ‘Image’ > ‘Adjust’ >‘Brightness/Contrast’. *(time 01:04, file S1)*
7. The sliders are moved to adjust the brightness and contrast manually.
8. The image parameters are changed by selecting ‘Image’ > ‘Properties’. *(time 01:33. File S1)*
9. The pixel measurements are changed to μm measurements, found in the image info in GDM.
10. The adjusted images are then saved in TIFF format by clicking ‘File’ > ‘Save as Image Sequence’. *(time 01:59, file S1)*

The following steps were then undertaken in each program to segment each of the image stacks.

*(1) Fiji/TrakEM2 (Image Analysis Fiji)*

1. The image stack to be analysed is opened in Fiji, as detailed above.
2. Then a new TrakEM2 file is created by selecting ‘File’ > ‘New’ > ‘TrakEM2 (Blank)’. *(see from time 00:04 in supplementary video file S2)*
3. Two new windows will open. One where new objects will be created and the other is the analysis window.
4. Images are imported in the analysis window by right clicking and selecting ‘Import’ >‘Import Stack’. *(time 00:14, file S2)*
5. User-defined objects within the data are created in the organiser by right clicking ‘Anything’ > ‘Add New Child’ > ‘Area_List’. *(time 01:10, file S2)*
6. Dragging ‘Anything’ into the middle column will create a new folder and then dragging ‘Area List’ will create a new object in that folder.
7. The object is renamed by right-clicking the ‘Area_List’ > ‘Rename’.
8. The object appears replicated in the ‘Z Space’ tab in the analysis window and has to be selected for segmentation. *(file 01:41, file S2)*
9. The brush tool is then used to draw an outline around the object to be segmented. The selected area is then filled in by holding the shift button and clicking in the centre of the outline. *(time 01:54, file S2)*
10. This is then repeated over each slice and for each user-defined object to be segmented.
11. Once the segmentation of defined object(s) is completed through the data stack, the results can be viewed in the 3D viewer. This is accomplished by right clicking in either window and selecting ‘Show in 3D’. *(02:54)*
12. The 3D reconstructions will appear in a new window.
13. The model is saved as an .obj file and can then be exported to other packages if required. *(time 04:57, file S2)*
14. 3D reconstructions created in Fiji were analysed in Blender.

*(2) Blender (Blender)*

1. The Neuromorph tools were installed via ‘Install from File’ > ‘File’ > ‘User Preferences’. *(see from time 00:24, in supplementary video file S3)*
2. The tools then appear in the side panel in the main interface under ‘Misc’. *(time 01:42, file S3)*
3. The .obj file is then opened by clicking on a tab in the bottom right hand panel marked ‘Scene’ > ‘Import Object’. *(time 02:21, file S3)*

*(3) Microscopy Image Browser, MIB (Image Analysis MIB)*

It can be opened through Matlab or as a standalone program, the information on how to do either is found on the MIB website.

1. The ‘Directory Contents’ panel is used to navigate to the location of the TIFF image stack to be imported. *(see from time 00:41in supplementary video file S4)*
2. All the images are highlighted, right-clicked and ‘Combine Selected Datasets’ is selected.
3. The image stack will open.
4. The image parameters are changed by going to ‘Dataset’ > ‘Parameters’. *(time 02:03, file S4)*
5. Then a model file is started by clicking ‘Create’. *(time 02:52, file S4)*
6. To add objects the ‘plus’ button is clicked and an object is added to the columns in the panel. *(time 03:01, file S4)*
7. A separate window will open where the name of the object can be changed and it will then appear in the column.
8. In MIB one object has to be segmented at a time, as pixels can only be selected and assigned to one object at a time.
9. The colour of the object can be changed by double-clicking on the colour square to the right of the object name.
10. In MIB there are a variety of methods that can be used to perform segmentation.

Manual

1. Clicking ‘Selection Type’ enables one to select a manual segmentation tool. The brush tool is a common one to use. *(time 03:37, fíle S4)*
2. The outline of each object can be drawn and repeated through each slice. Pressing Shift and F will fill in each object throughout all slices.

Interpolation

1. Interpolation was used to speed this process up by only drawing on every nth slice and then clicking ‘I’ or going to ‘Selection’ > ‘Interpolation’. *(time 04:22, file S4)*

Thresholding

1. Thresholding works by selecting pixels based on a user inputted contrast range.
2. There are two types of thresholding used B/W thresholding and the Magic Wand tool.
3. B/W thresholding will threshold the entire image and select pixels based on a user-inputted range, which is altered until the correct selection is made. *(time 05:16, file S4)*
4. This is applied to all slices within the stack by selecting ‘All’ for B/W thresholding
5. To use the magic wand tool, a defined range is set in advance and the radius of the tool is also changed. *(time 06:07, file S4)*
6. Then by clicking on a single pixel in the desired structure, pixels within the set range and tool radius will be selected.
7. By selecting the ‘3D option’ the process will be applied in some of the dataset, but not all, so if the structure is large this will have to be repeated.
8. The selection was then checked for any errors and adjusted manually using the brush tool, the tool can become an eraser by holding down the ctrl key.
9. The selected pixels, by whichever process, will appear green on the image.
10. These are assigned to the correct object by ensuring that the correct object is ticked in the segmentation panel and pressing Shift and A.
11. Once segmentation is finished (by whichever chosen process) the model file can be saved in a variety of formats.
12. For the duration of the segmentation the matlab file format (.mat) is preferred and the model will automatically save in this format. *(time 07:56, file S4)*

Saving for opening in Amira

1. To save the model file as a different file type go to ‘Model’>‘Save Model As’
2. Then in the file options select any of the ‘.am’ file types.
3. The new model file can now be opened in Amira (See Reconstruction under Amira)

*(4) Amira (Image Analysis Amira)*

1. Dataset is imported by selecting ‘Open data’, highlighting all the TIFF images and clicking open. *(see from time 00:05 in supplementary video file S5)*
2. A new window will open and under voxel measurements the pixel measurements, as mentioned at the beginning, are inputted. *(time 00:15, file S5)*
3. In the main interface a file will appear which corresponds to the image stack and a single orthoslice will automatically appear in the viewing area on the left.
4. Segmentation is started by right-clicking the main file and selecting ‘Edit New Label’. *(time 00:45, file S5)*
5. The view will switch from the ‘Project Panel’ to the ‘Segmentation Panel’.
6. In the segmentation panel new objects are added by clicking ‘Add’ and a new object will appear in the panel. *(time 00:58, file S5)*
7. Name can be edited by double-clicking the object and the colour changed by right clicking on it.
8. The bottom of the segmentation panel is where the segmentation tools are.

Manual and Interpolation

1. The brush symbol is for the manual tool, which is used to highlight objects. *(time 01:17, file S5)*
2. For interpolation every nth slice can be manually annotated and then interpolation clicked, found under the ‘Selection’ tab. *(time 02:16, file S5)*
3. Any object selection will appear coloured and has to be assigned to a label. This is highlighted in the top box by selecting the ‘plus’ sign.

Thresholding

1. The magic wand thresholding tool is selected by clicking on the magic wand symbol. *(time 03:21, file S5)*
2. When it is selected a graph will appear with sliders, which are adjustable
3. Using the tool a pixel in the structure to be segmented is selected, and a pointer will appear on the graph to show where on the contrast scale that pixel appears.
4. An area will then become highlighted and can be adjusted with the sliders until the right pixels are selected.
5. This can be done in 3D by ticking ‘3D’
6. In the segmentation panel the object can be viewed in 3D while segmenting by choosing the 4 panel viewer. *(time 05:18, file S5)*

Reconstruction

7. Back in the project panel a secondary main file will have been created which corresponds to the new objects created in the segmentation panel.
8. The model, all of the segmented objects, can be rendered by right clicking the labels file and selecting ‘Generate Surface’ > ‘Apply’. *(time 06:48, file S5)*
9. Another file will appear connected to the Labels file.
10. Right clicking this on ‘Surface’ and selecting ‘Surface View’ will render the 3D model in the viewing window.

## Results

Figure 2 illustrates a montage of serial-sectioned raw data SBF-SEM images from the psoas skeletal muscle (late fetus). Nine consecutive images are shown from a dataset of 93 serial images. This dataset was used to illustrate the application of the different image analysis programs and protocols. From the data the nucleus, nucleolus, chromatin and mitochondria were all segmented and reconstructed using the tools detailed in the methods. Further information on which tool was specifically used for each structure is included in the following results and figures.

**Figure 2.**
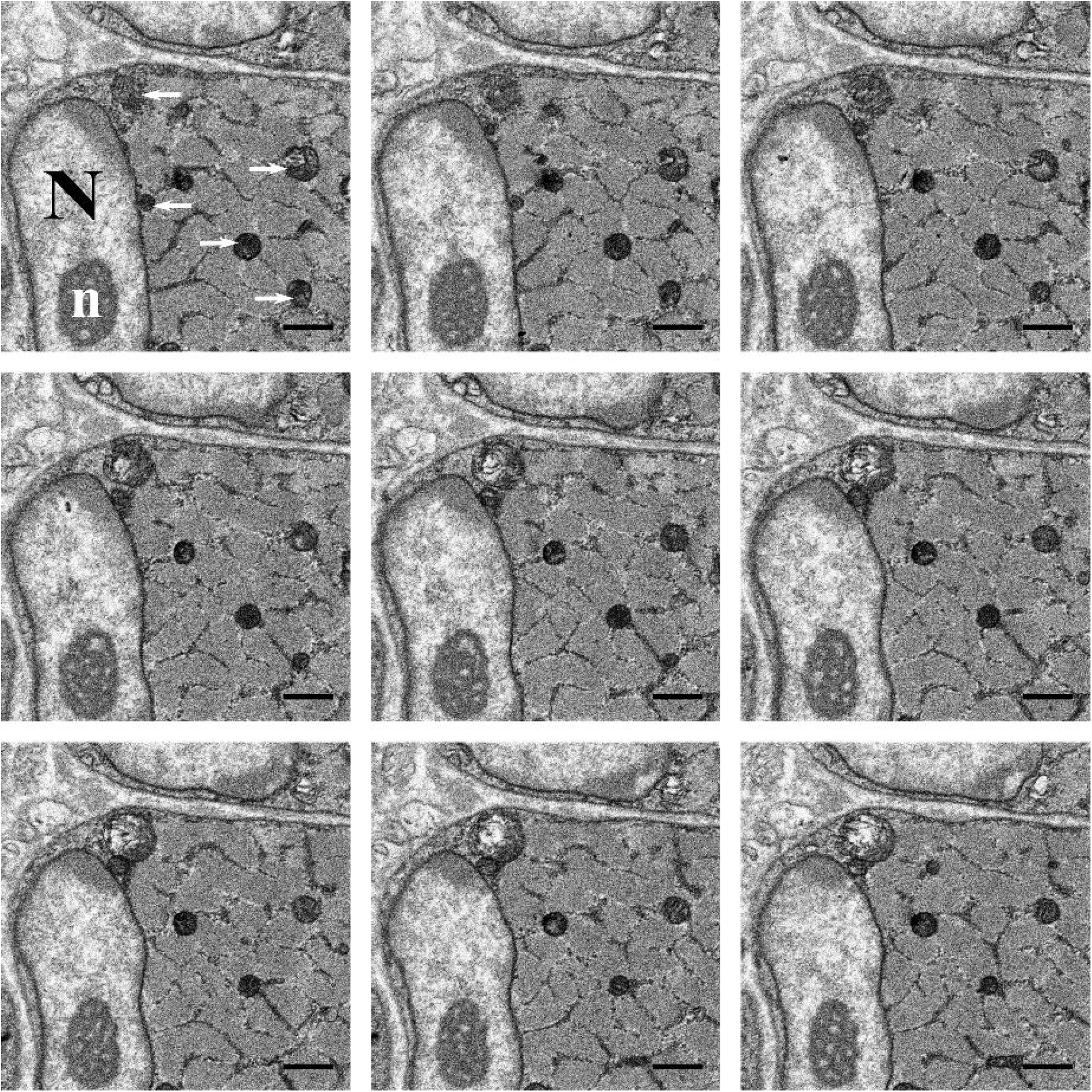
Example of SBF-SEM Image Series. 9 consecutive images (viewed from left to right) from a stack of 93 serial images of a portion of a muscle cell from the skeletal muscle psoas from a late fetus guinea pig. In the first image the nucleus can be seen, labelled with an ‘N’, as well as the nucleolus, white ‘n’, and the mitochondria, labelled with white arrows. Over each slice, of 70nm, the structures change shape, as shown in the images. All scale bars are 1 μm.

A timing analysis was also performed over 20 slices for each different method used to segment the nuclei and mitochondria. Interpolation was used to segment the nuclei in MIB and Amira, which was quicker than the manual segmentation performed in Fiji, with MIB being the fastest between all 3 programs. The thresholding method in Amira was the quickest method to segment the mitochondria. Overall the semi-automated tools, Amira specifically, segmented the structures fastest, when combining the nuclei and mitochondria timing analysis.

### Segmentation of Cellular Structures

The nucleus of the cell was segmented first. This was done manually in Fiji and with the use of interpolation in Amira and MIB. Visually the 3D models of the nuclei are similar and the volumes are also similar between each of the programs (Figure 3). Although using interpolation speeds up the segmentation process, errors did occur on the slices that were interpolated and correcting these errors adds to the time taken to segment. Further analysis was performed to test the accuracy of the interpolation in MIB, comparing the uncorrected interpolated nucleus to a manually segmented one (Figure 4). There is a differing level of detail between each of the models. Specifically the folds of the nuclear membrane are not as detailed in the interpolated models as in the manually segmented one. However the volume of the nuclei from the quantification analysis are similar, with only 1.98% difference and 1.37% difference between the corrected and the every 10th slice and 5th slice interpolation respectively. Therefore for bulk quantification analysis, corrections may not be required. If, however, finer details are required for qualitative analysis the interpolation errors will need to be corrected.

**Figure 3.**
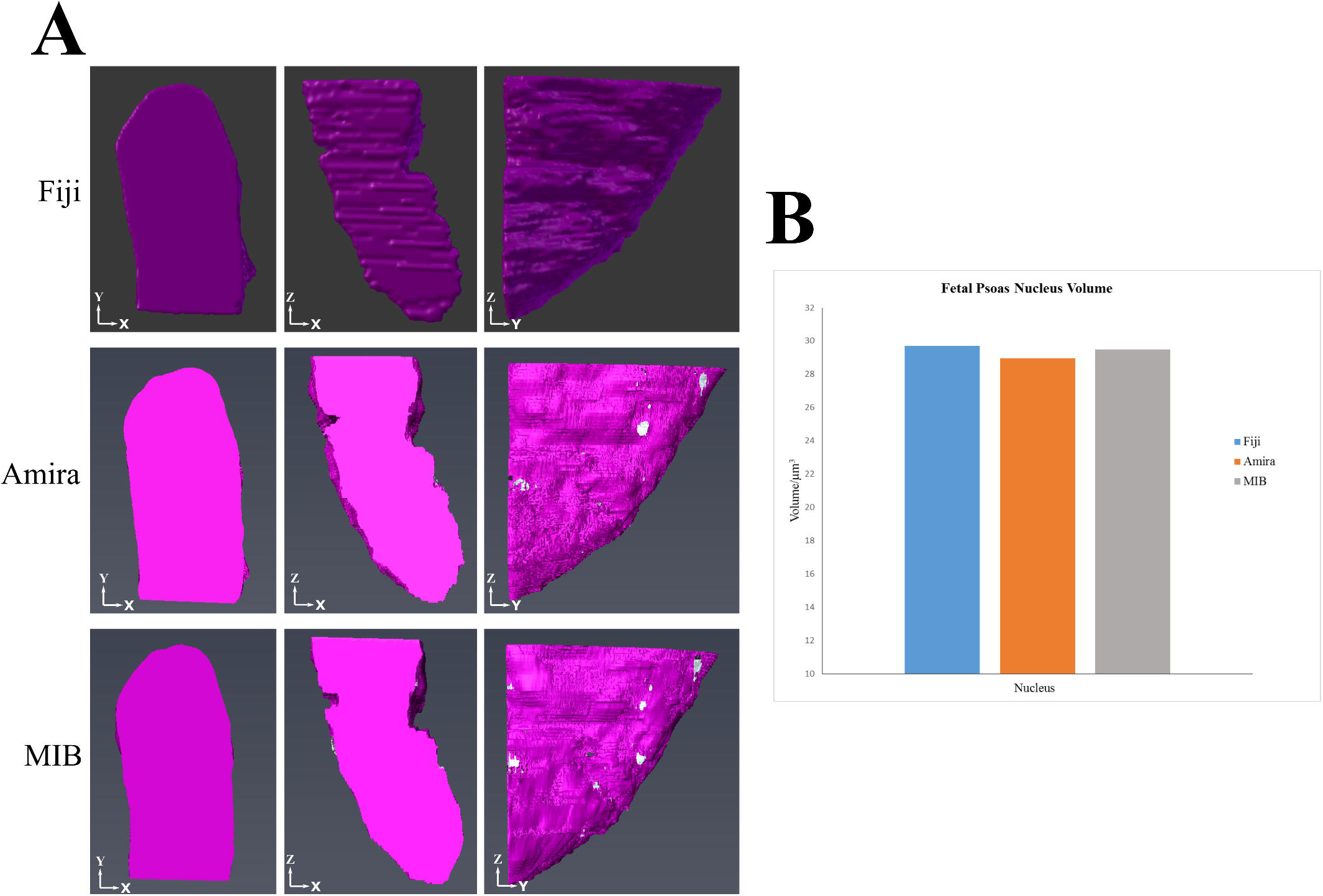
Examples of Digital Reconstruction of Nuclear Volume. (A) Reconstructions of the nucleus from fetal psoas at different orientations from the segmentations performed in Fiji, Amira and MIB. The reconstructions of the nuclei show no differences between the different programs. (B) Volume measurements of the nucleus from each of the programs, which again are similar between each of the programs used to reconstruct the nucleus.

**Figure 4.**
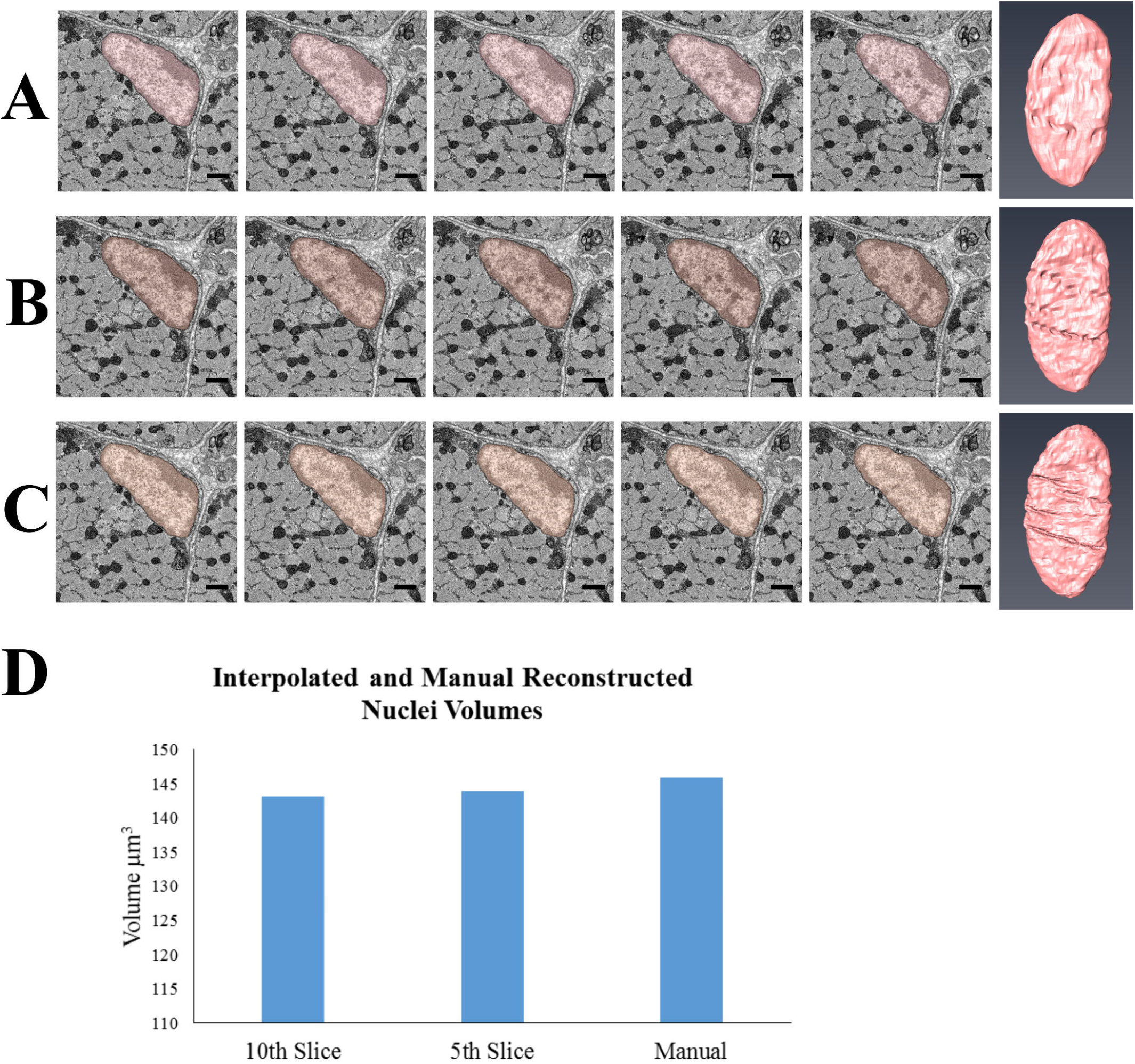
Analysis of Accuracy of Interpolation Method for Segmentation. (A), (B) and (C) each show 5 images with the nucleus segmented and the final reconstruction of the nucleus, from late fetal psoas all performed in MIB. (A) Shows the nucleus segmented using interpolation when every 10th slice has been manually segmented, (B) from every 5th slice and (C) is a nucleus which has been segmented manual. In the images there appear to be small differences between the segmentations, either the selection has not reached the boundary or goes over it. In the reconstructions the nuclear folds are not as detailed in (A) and (B) when compared to the manual reconstruction in (C). (D) Volumes from each of the segmentations. All Scale bars 1 μm.

After the nucleus, the darker and dense chromatin within the nucleus was segmented. The segmentation and the final reconstruction can be seen in Figure 5A. Thresholding was used in MIB and Amira to select the darker pixels that correspond to the chromatin. During the thresholding of the chromatin in both Amira and MIB there were some errors in the selection, for example incorrect selection of the nucleus boundary, and further user input was required to ensure that the correct selection was made. On the other hand, manual segmentation is subjective to the user and what they class as being part of the chromatin. Thresholding is one possible way in which this type of bias can be reduced.

**Figure 5.**
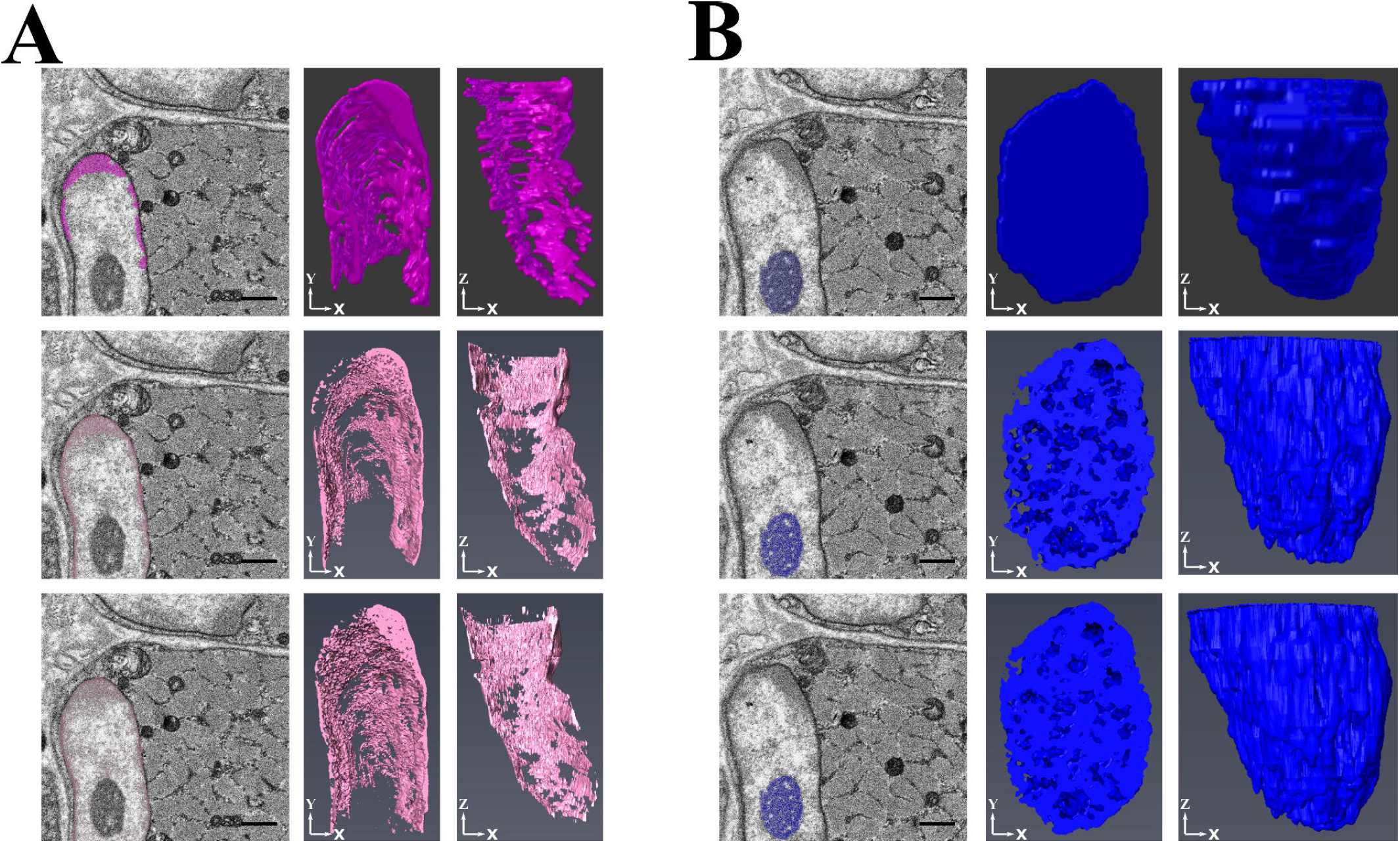
Examples of Reconstruction of Chromatin and Nucleoli. (A) 3 single snapshots from the data series with the chromatin segmented and the reconstructions at different orientations performed in each of the image programs, Fiji, Amira and MIB. (B) 3 single snapshots from the data series with the nucleolus segmented and reconstructions of the nucleolus from the 3 programs. The nucleoli reconstructed in Amira and MIB show more detail of the ‘web-like’ appearance of the nucleolus. All Scale Bars 1 μm.

The next to be segmented was the nucleolus. The segmentation has to be performed in this order due to the nature of the segmentation in MIB and Amira. As mentioned earlier, in these two programs pixels can only be assigned to a single structure. After the initial segmentation of the nucleus, all pixels are assigned to it. Then when the chromatin is segmented the thresholded pixels are reassigned from the nucleus to the chromatin. However, when the chromatin is thresholded the nucleolus is also selected, as the pixels are of a similar contrast. By segmenting the nucleolus after the chromatin, the pixels are reassigned to the nucleolus. This has to be kept in mind whenever segmenting a larger structure (nucleus) and the inner detail of it (chromatin and nucleoli) when using programs like Amira and MIB.

The segmented image and reconstruction of a nucleolus can be seen in Figure 5B. As seen in the raw images the nucleoli are a range of contrasts, with light and dark pixels. When thresholded, only the darker pixels are selected, which cause gaps in the model, whereas when it is manually segmented the entire nucleolus area is segmented. This is one example of when thresholding can be used to show intricate internal structures of an object.

Once the nuclei and associated structures were completed, the mitochondria were segmented. The decision was made not to interpolate the mitochondria due to their small size and complex nature, highlighted in Figure 6A and 6B. Instead the mitochondria were thresholded in Amira and MIB. When the 3 models are compared there are differences (Figure 6C). Mitochondria consist of a range of contrasts and in the raw images the detail of the cristae (appearing as dark inner membranes) can be seen. When thresholded only the darker pixels, the outer membrane and cristae, are selected. This gives the thresholded models, a broken appearance although the location and arrangement of the mitochondria can still be discerned and with further user input the appearance could be improved.

**Figure 6.**
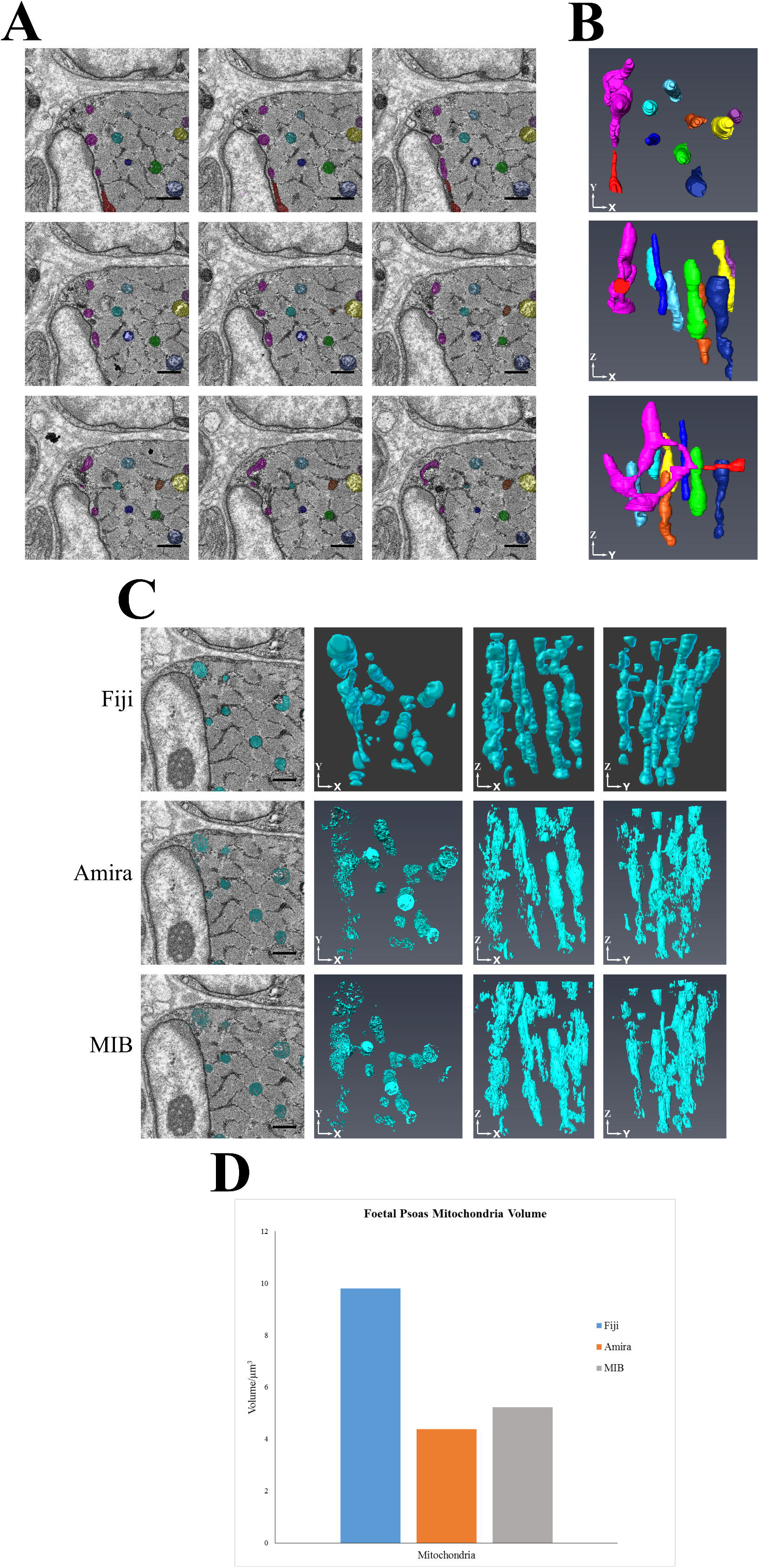
Examples Showing the Complex Morphology of Mitochondria. (A) 9 consecutive images from the late fetal psoas (viewed from left to right). The mitochondria have been segmented individually in different colours, in MIB, and their corresponding 3D reconstructions, from Amira, can be seen in (B). Showing how mitochondria change over each 70nm slice, requiring observation from the user to ensure that the correct structure is selected. All scale bars 1 μm. (C) 3 single snapshots from the data series with the mitochondria segmented and the reconstructions of the mitochondria at different orientations. The mitochondria, reconstructed in Amira and MIB, appear broken due to the selection method used. (D) Volume measurements of the mitochondria, which show that there is a large difference between the Fiji segmentation and the Amira and MIB segmentations, due to the broken appearance seen in the reconstructions. All Scale Bars 1μm.

A second dataset was taken at a higher resolution, higher magnification and thinner section thickness, specifically to reconstruct the mitochondria with greater detail. Figure 7 presents the results from the reconstruction of these datasets, showing that it is possible for the cristae of the mitochondria to be reconstructed in detail. However, importantly, quantitative analysis of the thresholded mitochondria results in a smaller volume than the manually segmented mitochondria, due to only the darker pixels being segmented. Therefore if quantification of mitochondrial volume is required manual method will have to be used to ensure correct measurements are made.

**Figure 7.**
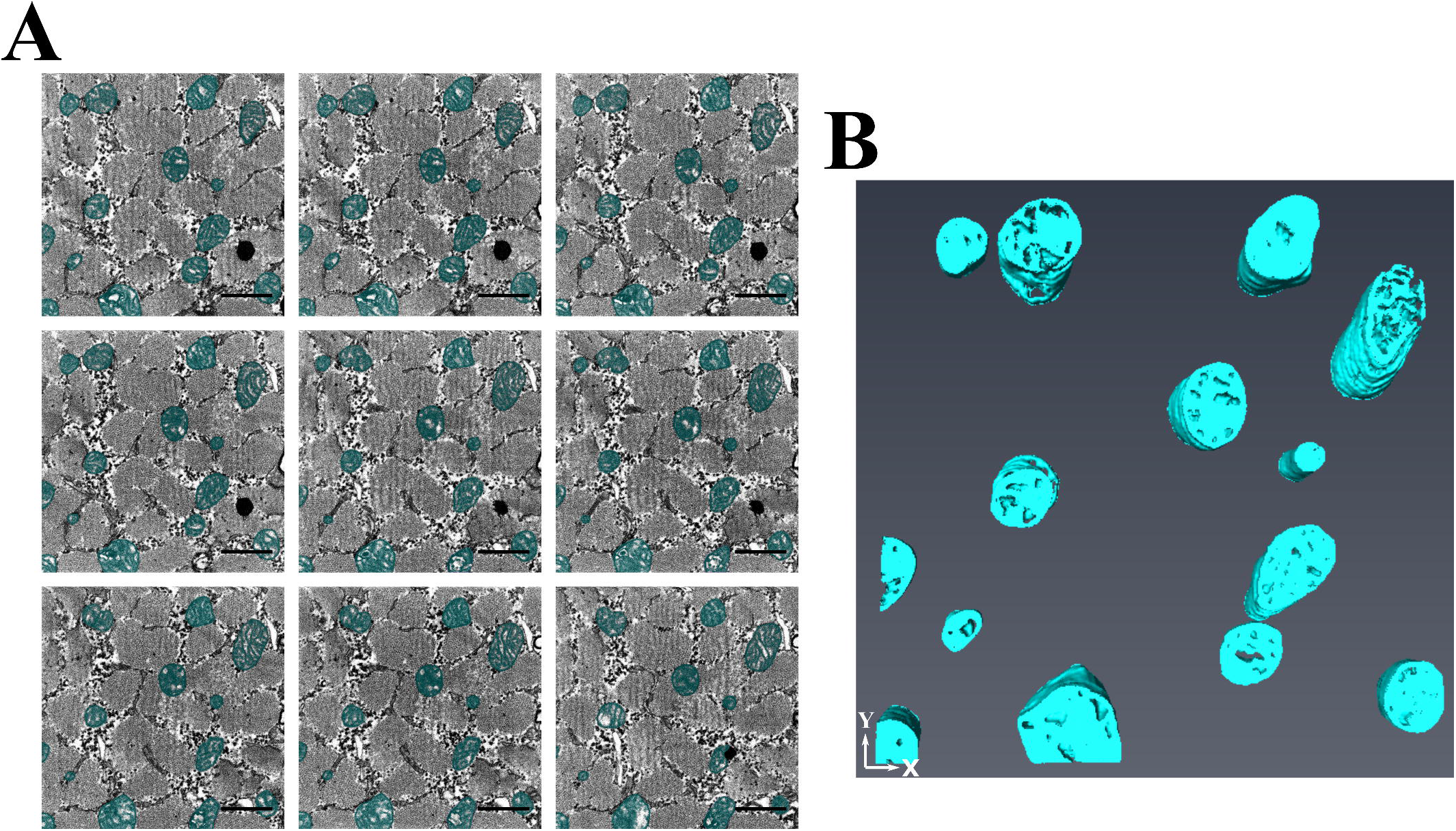
Digital reconstruction of mitochondria from a dataset with thinner sectioning and higher magnification. (A) Are 9 consecutive images (viewed from left to right) from a larger dataset from the fetal psoas muscle. The images were taken at high magnification (18.63kx), high resolution (5nm) and the block was sliced at 40nm. (B)The subsequent reconstructions from a portion of the total stack to highlight the reconstruction of the cristae of the mitochondria using the thresholding tool in MIB and Amira to reconstruct the mitochondria. Scale bar is 1 μm.

The final reconstructions of the skeletal muscle are shown in Figure 8. Although the general appearance of the models are similar in each of the programs, there are some differences caused by the methods used to segment the structures of interest. For example the mitochondria are more fragmented in the MIB and Amira models as they have been thresholded compared to the more dense structures seen with Fiji’s manual segmentation.

**Figure 8.**
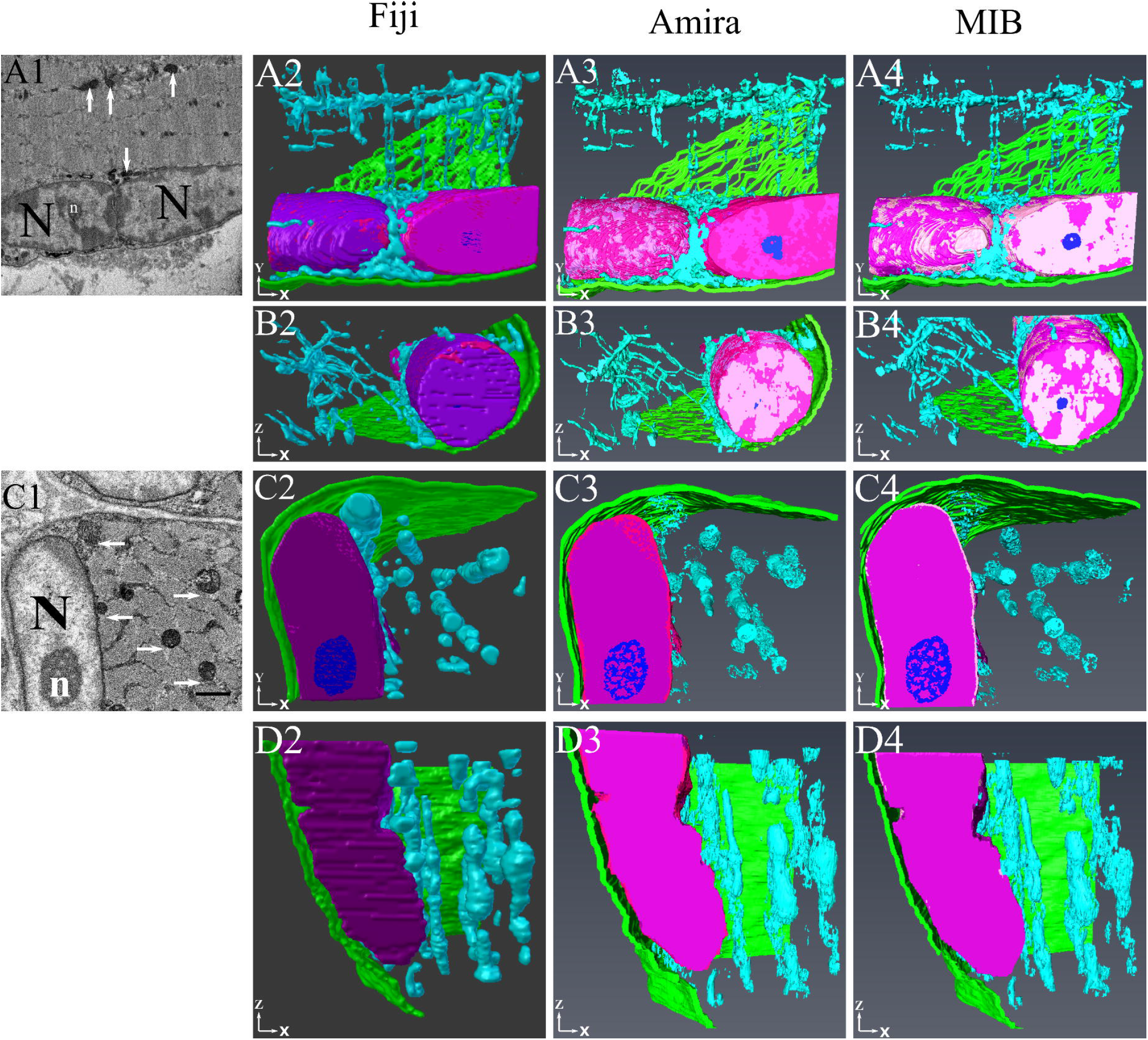
Examples of 3D Reconstructions of Segmented Structures. This diagram depicts examples of the assembled reconstructions of all segmented features from two separate skeletal muscle SBF-SEM datasets. Rows A and B show results from adult soleus muscle (from X serial sections; panel A1 indicates a snapshot SBF-SEM image). Rows C and D show results from fetal psoas muscle (from X serial sections; panel B1 indicates a snapshot SBF-SEM image). Rows (C) and (D) The following features are colour-coded in the reconstructions: mitochondria, light blue; nuclei, dark pink/purple; chromatin, light pink; nucleoli, dark blue; plasmalemma, green. All scale bars 1μm.

## Discussion

Serial block face SEM is a powerful tool for cellular examination and, when combined with image analysis software, can provide detailed qualitative and quantitative data. However, getting started with the software is not easy for new researchers and there is a risk they will under utilise their data. Here we have explained the terminology of many analytical features found in the programs and provided step-by-step protocols to instruct users.

We have compared manual and semi-automated segmentation methods in order to help researchers chose the best options for their analysis. The results show that the semiautomated methods are less time consuming but are not always accurate, as shown by the quantification results of the mitochondria segmentations. However this segmentation was performed with the thresholding tools, which work best on structures of a single contrast that are distinct from their surroundings. Thresholding can be a useful way to highlight the finer details of structures, such as the cristae of the mitochondria, the web-like appearance of the nucleolus and the chromatin within the nucleus. In all cases these semi-automated tools require some form of manual input and manual correction. A prime example of this is when using interpolation, which is best suited to larger structures that do not change much over each slice, like the nucleus of a cell, a whole cell or a large portion of the tissue. To optimise results, each object of interest should be assessed individually in terms of the segmentation method. Using a range of tools to efficiently and accurately segment multiple structures in a sample will give better results than trying to use a ‘one-method-fits-all’ approach.

Although the results shown are derived from guinea pig skeletal muscle, the described approaches have been repeated on other types of tissue (guinea pig cardiac muscle and locust optic lobe), which validate the findings shown. From the results a workflow has been devised, Figure 9, to aid researchers new to image analysis. It provides recommendations for segmentation tools based on a variety of factors, such as the contrast and size of the structure, presence of any membrane boundaries, the size of the dataset and most importantly the objective of the analysis. The result can be qualitative, quantitative or both and knowing which is required prior to analysis is important, as shown by the results in this paper.

**Figure 9.**
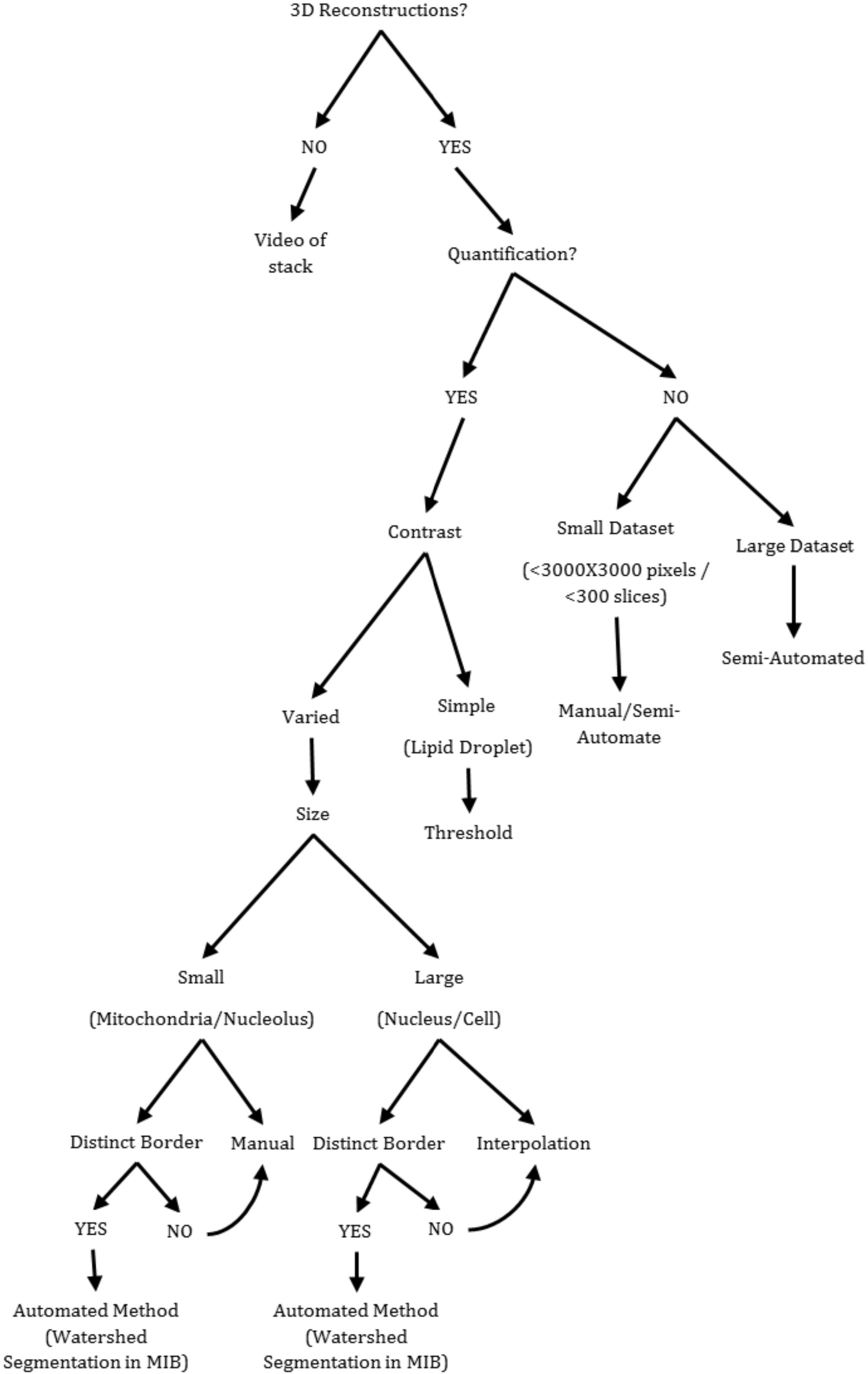
Proposed workflow to aid in decision making when choosing appropriate segmentation methods for analysis of SBF-SEM data. The majority of these segmentation methods can be used in MIB and Amira, with some exceptions, the watershed segmentation which is not shown in this paper. The decision to use either MIB or Amira to perform the segmentations will depend on user preference and access to the software, as previously shown the two programs yield similar results.

Knowing the objective is not only important for the image analysis but also prior to that, when the data sets are collected. For example, if low-resolution images are collected to reconstruct a whole cell, it is no good deciding after collection that the organelles should also be reconstructed, as they will not have the clarity. The resolution of the image collection should be determined by the smallest structure likely to be of interest. However, although decisions on how to collect data in pursuit of biological questions are made at the time of SBF-SEM scanning, the outcomes are only revealed upon viewing the serially collected digital images. Indeed, this is one of the key benefits of SBF-SEM. Therefore, if tissue is plentiful, an iterative approach to SBF-SEM data collection and analysis can be beneficial in revealing much new biological information from complex cell/tissue structures.

By adjusting the data collection parameters - section thickness, magnification, number of pixels - the resulting images will have different resolution, thus altering the appearance of the final reconstructions. A thinner section thickness will result in a more detailed reconstruction, as smaller changes in structures will be imaged. A higher resolution results in a clearer image, so a better distinction between cellular structures and higher detail of the structures is achieved. The resolution, magnification and number of pixels are all linked. A higher magnification results in a higher resolution image but this decreases the field of view. To maintain the same field of view but still increase resolution, the number of pixels can be increased. For example, to achieve x nm resolution an image that is 1000 by 1000 pixels will have to have a higher magnification than an image that is 3000 by 3000 pixels. However an increase in pixel number also increases the resulting image file size, which can be difficult to handle without a high-powered computer. In addition, if the time taken to collect an image is very high, the electron dose can have a detrimental effect on the resin and cause subsequent sectioning artefact. An alternative would be multiple regions of interest (ROIs) at a higher magnification but lower pixel number. However ROIs that overlap might be affected by the increased exposure to the electron beam, as the areas are scanned repeatedly. Thus some compromise may be required to balance the need for high-resolution images but also images free from sectioning and imaging artefacts.

We have provided a workflow to recommend segmentation tools and also a detailed step-by-step protocol for 4 programs, one of which is newly developed (MIB). The aim is to provide a resource for researchers to refer to when starting their analysis. Although there have been several publications using SBFSEM, the detail about the segmentation methods is limited. We carried out a survey of the recent literature (published between 2013 and 2017) where SBFSEM has been utilised and found that although all stated which program they had used, only around half stated the specific tools utilised. Of these, only a small number of the articles described the segmentation process in detail, with the majority simply stating the type of segmentation. The more detailed papers tended to be more focused on the methods whereas the papers with fewer details tended to be more biological and used SBFSEM in conjunction with other techniques. A detailed analysis of the different software and their functionalities can be found in Kittelmann et al (2016). Earlier publications were reviewed by Borrett and Hughes (2016) and they also noted that there is often a lack of information given in articles on the methods used to analyse data from SBFSEM.

From the literature survey it was also apparent that different terms are used to describe the segmentation tools, for example manual segmentation was sometimes referred to ‘by hand’ and thresholding as ‘intensity-based’ or ‘contrast-based selection’. This could lead to confusion for those new to this type of analysis. Since its release in 2004 the use SBFSEM has steadily grown. In 2016 there were 39 articles published and so far in 2017 (April) there have been 13 publications using this technique. With this burgeoning interest there has been an increased demand for training in the analysis of data. This has drawn attention to the need for clarity and consistency in the reporting of methods of data analysis and interpretation, both qualitative and quantitative.

In addition, we recognise that this is a fluid research environment where computational advances are rapid. Therefore, our provision of openly accessible raw datasets enables researchers who are developing new analysis approaches to compare the functionality of new tools with those used here. All researchers should be encouraged to deposit future SBF-SEM datatsets at https://www.ebi.ac.uk/pdbe/emdb/empiar/, or similar open-access repositories.

We have not only provided detailed step-by-step protocols but also videos to run alongside these protocols. We have found that when teaching others to use new programs, it is much easier for them to learn if they can see the software layout and which buttons to press rather than relying purely on a description.

In conclusion, SBFSEM is a powerful tool for analysing cellular structures with high resolution in x-y-z planes. However current publications do not always give enough information on this process and with the myriad of programs and tools available it can be a daunting task for new researchers to train themselves. However by following a logical workflow it is possible to obtain qualitative and quantitative data on multiple structures from a single dataset, maximising output from valuable tissue.

## Acknowledgements

*The Research was funded by BBSRC grant BB/MO12093/1*

Raw data can be accessed from http://www.ebi.ac.uk/pdbe/emdb/empiar/ with Accession code: EMPIAR-10092

